# A Dynamic Bottom-Up Saliency Detection Method for Still Images

**DOI:** 10.1101/2022.03.09.483582

**Authors:** Leila Sadeghi, Shiva Kamkar, Hamid Abrishami Moghaddam

## Abstract

**Introduction:** Existing saliency detection algorithms in the literature have ignored the importance of time. They create a static saliency map for the whole recording time. However, bottom-up and top-down attention continuously compete and the salient regions change through time. In this paper, we propose an unsupervised algorithm to predict the dynamic evolution of bottom-up saliency in images.

**Method:** We compute the variation of low-level features within non-overlapping patches of the input image. A patch with higher variation is considered more salient. We use a threshold to ignore less salient parts and create a map. A weighted sum of this map and its center of mass is calculated to provide the saliency map. The threshold and weights are set dynamically. We use the MIT1003 and DOVES datasets for evaluation and break the recording to multiple 100ms or 500ms-time intervals. A separate ground-truth is created for each interval. Then, the predicted dynamic saliency map is compared to the ground-truth using Normalized Scanpath Saliency, Kullback-Leibler divergence, Similarity, and Linear Correlation Coefficient metrics.

**Results:** The proposed method outperformed the competitors on DOVES dataset. It also had an acceptable performance on MIT1003 especially within 0-400ms after stimulus onset.

**Conclusion:** This dynamic algorithm can predict an image’s salient regions better than the static methods as saliency detection is inherently a dynamic process. This method is biologically-plausible and in-line with the recent findings of the creation of a bottom-up saliency map in the primary visual cortex or superior colliculus.

## 1. Introduction

The Visual system has a critical role in our daily life. When we face a new scene, our brain is unable to process all data immediately. Therefore, cognitive factors and neural processes are used to analyse some parts of data in order. First, bottom-up attention involves. It finds the most salient area of the scene using the low-level features already extracted in the retina and visual cortex [1, 2]. Nowadays, we know that this stimulus-driven mechanism is a built-in compartment and no training is required for saliency detection using bottom-up visual attention [3, 4]. Indeed, the human visual system has evolved to ignore redundant information and guide eye movements to distinct areas quickly and involuntarily [5]. As time goes by, the extracted information travel to other areas of the brain and feedback from memory is received. This is when top-down attention overrules [6]. These two types of attention compete continuously and make the subject fixcate at different areas of the scene to get more visual data. Thus, the salient region-the most important parts of the scene fixated at by the subject-is continuously changing [7, 8]. However, it is common in the literature to accumulate all subjects’ fixation during the entire free-viewing task and create a single saliency map for the image stimulus. This way, time is ignored and the map does not show which parts were salient recently and which ones earlier. In this paper, we suggest developing a dynamic saliency detection algorithm to find the salient parts of an image through time.

The proposed method is inspired by the theory of a saliency map in primary visual cortex [1]. According to this theory, a bottom-up saliency map is created in V1 in a very short time immediately after stimulus onset [1], before the arrival and interference of top-down signals. In addition, it presumes that a single saliency map is created in V1 which cannot be considered as a simple combination of multiple feature maps [1, 2, 5]. In fact, firing rates of V1’s output neurons are higher in more salient regions of the image regardless of features.

Obviously, the proposed method must be evaluated using bottom-up ground truth-a fixation map that is generated in a short time interval after stimulus onset. Controversially, most of the bottom-up saliency detection counterparts have been evaluated on top-down datasets. Because their ground truths were generated using eye movements registered within multiple seconds after stimulus onset. However, this time is long enough to involve top-down attention as well as bottom-up. Time is the critical point to judge whether the eye movements are only based on bottom-up attention or top-down attention also plays a role. Discriminating these two types of attention is important mainly because they differ structurally. Bottom-up attention is stimulus-driven and does not need to be trained, while top-down attention is goal-driven and learning-based. So, we suggest using different strategies to model each. To evaluate properly, we break recorded eye movements into multiple intervals and create a saliency map for each separately. This way, a dynamic ground truth is generated that shows how saliency changes through time. We used MIT1003 and DOVES datasets for evaluation of the suggested algorithm.

The rest of the paper is organized as follows. In Section 2, some related works are reviewed. The details of the proposed algorithm are explained in Section 3. Section 4 describes the details of generating dynamic ground-truth for MIT1003 and DOVES datasets. Evaluation results are given in Section 5. Finally, Section 6 concludes the paper.

## 2. Related work

Basic saliency detection algorithms were computational models that have mimicked the brain function according to what neuroscientists and cognitive scientists reported. The first brain-inspired saliency detection algorithm was introduced in 1998 by Itti et al. [9] based on the feature integration theory [4, 10] and selective visual attention mechanism [11, 12]. In this method, the input image is scaled into nine spatial scales using Gaussian pyramids. Then, feature channels corresponding to color, intensity, and orientation are extracted from each scaled image. Subsequently, each channel is processed separately by applying center-surround interactions, normalization function, and aggregation across scales. Finally, feature maps for each channel are normalized and linearly combined to produce a master saliency map. This unsupervised framework became a starting point for modeling saliency detection. Later studies modified its building blocks by introducing new feature channels such as motion and flicker contrasts [13] or applying a different method for combining the maps [14, 15, 16, 17] such as non-linear combination [15, 18] or weighted aggregation [19]. Achanta et al. [20] demonstrated that a saliency map can be created by obtaining the filtered responses of the image in the CIE L*a*b* color space and then calculating the center-surround difference maps for each feature and their weighted composition. Several studies tried to reduce the computational complexity by adjusting the number of scales according to the size of the input image [21, 22].

Some researches are not completely in-line with the exact function of the brain and consider only some key aspects. For example, the entropy of the scene is considered as the main factor for processing visual features. A model is proposed in [23] for saliency detection based on information maximization theory. This method produces a saliency map using Shannon’s measure of self-information. In [22], a graph-based visual saliency (GBVS) model is proposed. This bottom-up saliency detection method introduces a novel graph-based normalization scheme and utilizes the Markovian approach. In GBVS model, activation maps on feature channels are produced, and after normalization, are combined with other maps. Likewise in [24], a graph-based method is proposed in which salient regions are detected by identification of the foreground and background cue based on color differences in LAB space. Torralba [25] and Oliva [26] provided a Bayesian framework to combine sensory information and prior knowledge via Bayes’ rules. Some models process the image in terms of pixels, and others use local and global features of the image for saliency detection. The authors in [27] benefited from the former and used color spatial information to estimate salient areas. Tie et al. [28] benefited from local, regional, and global features such as histogram, contrast, and spatial color distribution for saliency detection. In [29], covariance matrices of image features are used in addition to the simple color, intensity, and orientation features. In this model, the saliency of a local image patch is computed based on the distances between its covariance descriptor and those of the surrounding patches. It makes the feature integration process nonlinear. In [30], both local and global features are used to detect important parts of the scene by measuring the similarity between each image patch and other image patches. In [31], the double color opponency mechanism in V1, skin color detection, and background map are utilized for saliency detection. In [32], a regional contrast-based saliency detection method is proposed which simultaneously evaluates global contrast differences and spatial coherence. In [33], a bi-directional propagation model (BIP) is presented that utilizes the foreground and background propagation to select salient regions. In [34], a saliency detection model is proposed based on the spatial position prior of attractive objects and sparse background features.

Some approaches use neural networks and deep learning techniques such as convolution neural networks [35, 36, 37], sparse deep learning networks [38] Boltzmann machine [39], and ensemble deep neural network [40] for modeling bottom-up attention. In [41], a multiple convolution layers model is proposed to predict eye fixation which uses the end-to-end encoder-decoder network. For the image encoder, VGG16 architecture is used. This model is evaluated using, AUC, CC, NSS, and SIM measures on CAT2000, MIT1003 and SALICON datasets. In [42], a scanpath model is presented that utilizes the Long Short-Term Memory (LSTM) to generate the sequences of fixation. Also, during recent years, scientists utilized transfer learning. DeepGaze I [43] was the first method that used transfer learning for the saliency prediction. In [43], a new model of fixation selection is presented by using a pretrained deep neural network on the task of object recognition from Krizhevsky et al [44]. After DeepGaze I, transfer learning is used often based on ImageNet. In DeepGaze II [45], the fine-tuning of the VGG-19 features is used for saliency detection. The performance of saliency model is enhanced by using ResNet50 features instead of the VGG19 of DeepGaze II in [46]. In addition to the aforementioned algorithms which are mostly in the spatial domain, some models work in the frequency domain and benefit from Fourier transform, discrete cosine transform, etc. [47, 48]. In [49], Jian et al. introduced a wavelet-based salient patch detector to extract visual information for detecting image salient regions from separate color channels and intensity.

In contrast to the most existing methods, our proposed algorithm is dynamic and predicts bottom-up saliency through time. It is inspired by the theory of a saliency map in the primary visual cortex. The following section describes the method in detail.

## 3. The proposed dynamic bottom-up saliency detection algorithm

Humans and animals learn many things in an unsupervised manner [4]. Finding the salient parts of the scene using bottom-up attention is one of them [3]. We propose an unsupervised feed-forward saliency detection algorithm. The algorithm is inspired by the theory of “a saliency map in primary visual cortex” proposed in [1]. According to this theory, when the visual input emerges, the saliency map is generated immediately in the primary visual cortex and the eyes make a saccade to the most salient part. Top-down attention does not play a role in this short time just after presenting the stimuli. The bottom-up saliency map is a single map. No separate feature maps are generated or combined to create the final saliency map. Since this map is generated using conventional V1 cells’ firing rates. This output for saliency is universal: two neurons may have similar firing rates. However, they are selective to different features. It means that the saliency of their corresponding inputs is the same [1, 2]. In accordance with the above processing scheme, the proposed method relies on the variation of low-level features of color and orientation within small windows. The flowchart of the proposed method is shown in Fig. 1 and the details are given in the following subsections.

**Figure 1:**
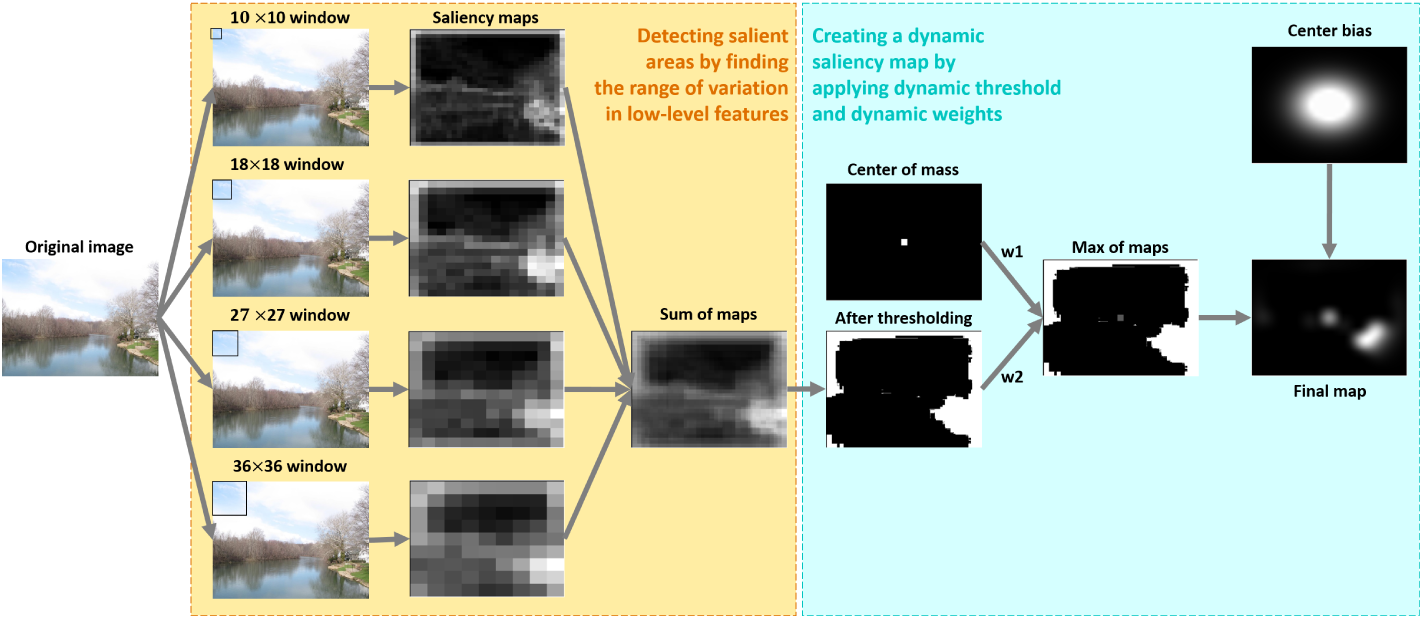
Flowchart of the proposed method

### 3.1. The core of saliency detection algorithm

Just after receiving the input image, three features including color, intensity, and orientation are extracted. We extract color information in CIE L*a*b color space. The intensity feature is extracted by averaging red, green, and blue channels in RGB color space. Orientation information is obtained by convolution of intensity image using Gabor filters. Then, all feature values are normalized within the range of [−1, 1].

These features were selected according to the physiology of the human visual system. It is commonly known that the retina is responsible for extracting color information thanks to the photoreceptors even before the information is transmitted to the visual cortex [2]. Also, V1 neurons can detect the edges with different orientations [1, 2]. The Gabor filter is linear and local and is proved to be a powerful tool for texture analysis due to its high resolution in both space and frequency domains. There are some evidence that simple cells in mammalian V1 can be modeled by Gabor functions [50, 51]. Accordingly, we convolve the intensity image with the Gabor filter (Eq. (1a)) to find the edges along four orientations of 0, 45, 90, and 135 degrees:

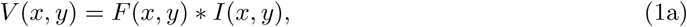

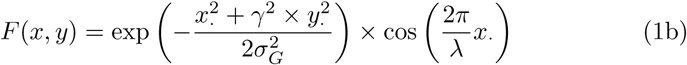

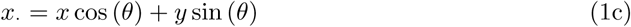

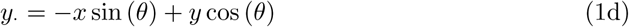

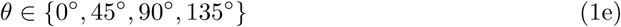

where *I*(*x, y*) is the input intensity image and ∗ denotes the convolution operation. *F* (*x, y*) is the Gabor filter function in the coordinates (*x, y*), and (*x*_·_, *y*_·_) is the polar coordinates of (*x, y*). The parameters of *γ, σ*_*G*_, *λ*, and *θ* are the spatial aspect ratio, standard deviation, wavelength, and orientation, respectively. The size of the Gabor filters is the same as the window size.

Features in different parts of the input image are processed within windows. It resembles the concept of receptive field (RF). The cells in the retina have very small RFs [52, 53]. This is also true for the neurons in the primary visual cortex. The overlap of these RFs is very small in V1 and increases in the upper layers of the visual cortex [2]. V1 neurons’ RFs are different in size. They vary approximately from 0.25 to 2 degrees of visual angle [53]. Retina’s cell’s RFs in fovea are smaller than 1 degree of visual angle [54]. Therefore, We considered four different window sizes of 10 × 10, 18 × 18, 27 × 27, and 36 × 36 pixels to cover the common RF size range between retina and V1. The sizes of 10, 18, 27, and 36 pixels correspond to 0.26, 0.47, 0.71, and 0.95 degrees of visual angle according to the data recording settings in MIT1003 [55].

Windowing helps us to decrease the dimension and still preserve important information. Using a window size of 10 × 10, a 500 × 500 image is converted to a 50 × 50 matrix. Each element of this matrix corresponds to a window and is determined by processing the information within that window. The element’s value is assigned by taking the summation of the range of each extracted feature (L, a, and b color features and four orientation features.) The range of a feature is defined as the difference between the lowest and highest values within the window. The summation operator takes the weighted sum of the range values. All features have the same weight of 1, except color feature b with the weight 0.2. This resembles the lower density of blue con cells relative to the green and red ones in the retina. When the values of all elements in the matrix are determined, we see that the minimum and maximum values of the matrix correspond to the two windows with the lowest and highest variability. It is widely known that the cells in retina and V1 have center-surround RF and encode the existence of a feature somewhere that its surround lacks it. We interpret it as a variation detector and use range as a variation measure.

After replacing each window with the corresponding sum value, a matrix is created. The maximum element of this matrix shows where the maximum variation exists. Therefore, the matrix values are comparable and encode bottom-up saliency regardless of features. To get a saliency map in the same size of the original image, we replace each element of the matrix with a 10 × 10 window. The value of the matrix’ element is copied to all cells of the corresponding window. The above process is repeated for different window sizes and finally we get 4 saliency maps with different resolution. The final saliency map is generated by summation of all 4 maps. In bottom-up attention, people look at the parts with the highest variability in terms of low-level features. Similarly, in the final saliency map, the maximum value shows the most variability and the most salient part in terms of low-level features.

### 3.2. Creating a dynamic saliency map

Sometimes, people look at somewhere near all salient parts. For example, before the subject decides to fixate on which part, or while making saccades from a salient area to another. Accordingly, we find the center of mass as a point that is somewhere between all detected salient areas by taking the average of all nonzero values in the saliency map. This strategy is used in a short time after stimulus onset. As the time goes by, the subject tends to fixate exactly on the salient area. After that, the less salient areas attract attention too. Equivalently, we set a variable threshold and remove values less than that in the saliency map. At the beginning, the threshold is high (0.95 of the maximum value in saliency map) and only the most salient parts are involved in creating the final saliency map. The threshold decreases 0.05 for each interval and becomes 0.9, and 0.85 corresponding to 100-200 and 200-300 time intervals. After applying the threshold, we merge the weighted versions of the saliency map and the center of mass using the maximum function. The weights are set dynamically; center of mass’s weight (w1) decreases through time and the weight for saliency map (w2) increases. The former (w1) follows an exponential function with the parameter 0.9. The later is calculated as w2=1-w1. Then, a pixelwise maximum function is applied on the weighted versions.

### 3.3. Center bias

When observers view an image on the screen, their tend to fixate on the central area, because it is assumed that the salient parts are biased towards the image center [56]. Also, when a person looks at a scene, small saccades along the horizontal direction are more likely to occur than the vertical direction [57]. The probability distribution of these saccades can be modeled as a Gaussian distribution. Accordingly, we define a 2D Gaussian with the standard deviation *σ*_*c*_ = 5 within a 33 × 33 center map. Then, it is resized to the size of the saliency map and is multiplied by it to enhance the central parts and fade the boundary (Eq. (2)):

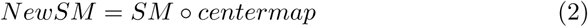

where *SM* , *centermap* and *NewSM* are the saliency map, the center map, and the final saliency map, respectively. The symbol ◦ shows hadamard product that is a pixelwise multiplication of two images. Finally, a 150 × 150 Gaussian Kernel with standard deviation 40 pixels smoothes the saliency map.

## 4. Datasets

Saliency maps generated by computational models should ideally select the regions which attracted the human visual attention. Therefore, to evaluate the proposed model, the data which are obtained by tracking eye movements are used. Various saliency datasets are available such as CAT2000[58], SALICON[59], TORONTO[60], DUT-OMRON[61], HDREYE[62], PASCAL-S[63], EMOd[64] and Le Meur[65] dataset. However, we could not benefit from their data mainly because only fixation lists were available and it was not clear when each fixation happens. In two datasets MIT1003[55] and DOVES[66], complete raw eye data with acceptable temporal resolution are released. Therefore, we used them for evaluation. MIT1003 contains 1003 images and their corresponding eye data for 15 viewers during 3-second free-viewing task ^1^. We broke this time to 14 time intervals: 0-100ms, 100-200ms, 200-300ms, 300-400ms, 400-500ms, 500-600ms, 600-700ms, 700-800ms, 800-900ms, 900-1000ms, 1000-1500ms, 1500-2000ms, 2000-2500ms, 2500-3000ms. We considered at least 100ms time intervals to make sure that there are enough fixation data to generate a ground truth. DOVES contains 101 grayscale natural images observed by 29 humans for 5-second. We broke this time to 18 time intervals: 0-100ms, 100-200ms, 200-300ms, 300-400ms, 400-500ms, 500-600ms, 600-700ms, 700-800ms, 800-900ms, 900-1000ms, 1000-1500ms, 1500-2000ms, 2000-2500ms, 2500-3000ms, 3000-3500ms, 3500-4000ms, 4000-4500ms, 4500-5000ms. For each time interval, the corresponding ground truth was generated using eyedata recorded in that interval. Figure 2 shows how this dynamic ground truth varies through time for a sample image from each dataset. This is in contrast to the common strategy of accumulating all data and generating a single ground truth for an image.

**Figure 2:**
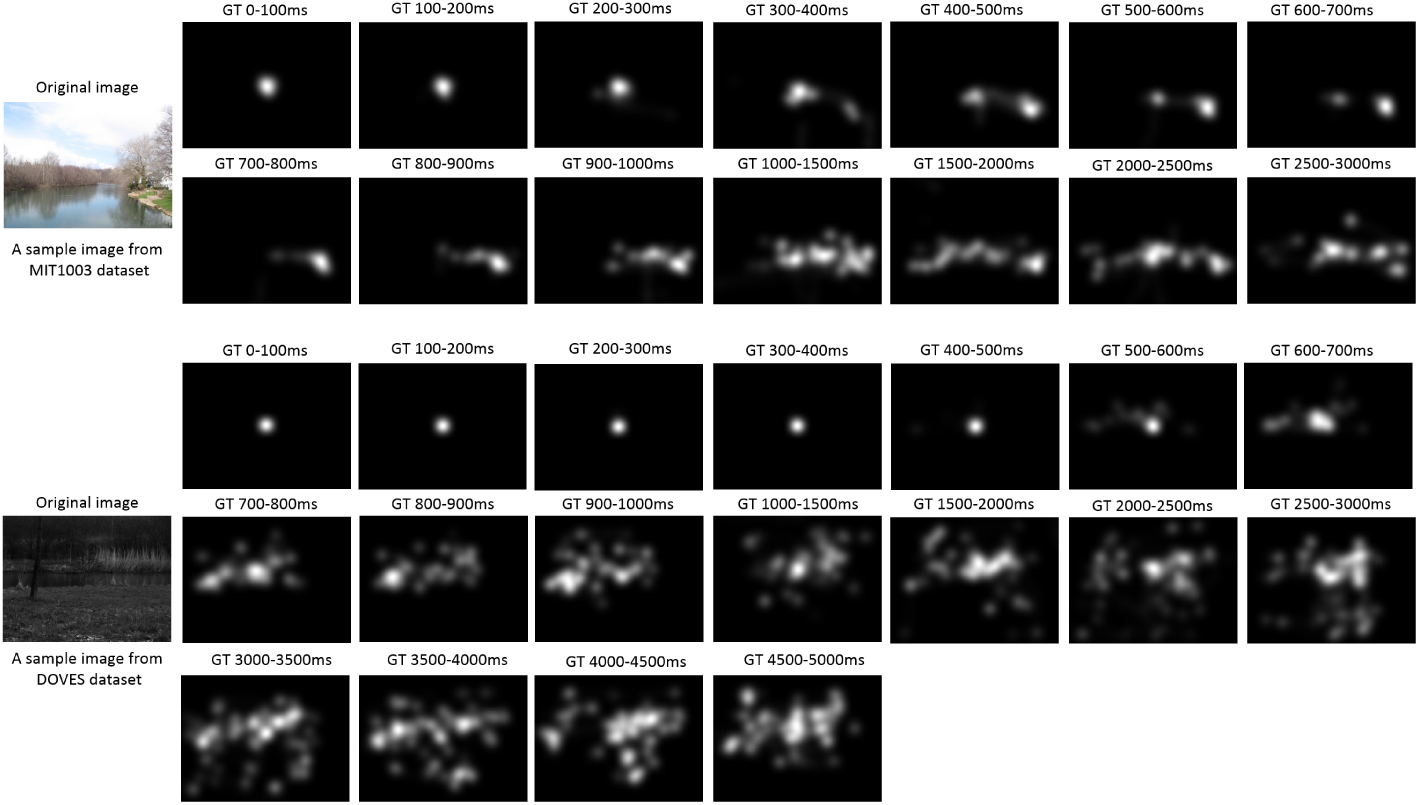
Dynamic ground truth for a sample image in MIT1003 dataset (top) and a sample image in DOVES dataset (bottom).

## 5. Evaluation

The algorithm was implemented using MATLAB R2018a on a standard PC with a 3.4GHZ Intel core i7 CPU and 16GB of RAM. Figure 3 shows the output saliency map for a sample image in both MIT1003 and DOVES dataset.

**Figure 3:**
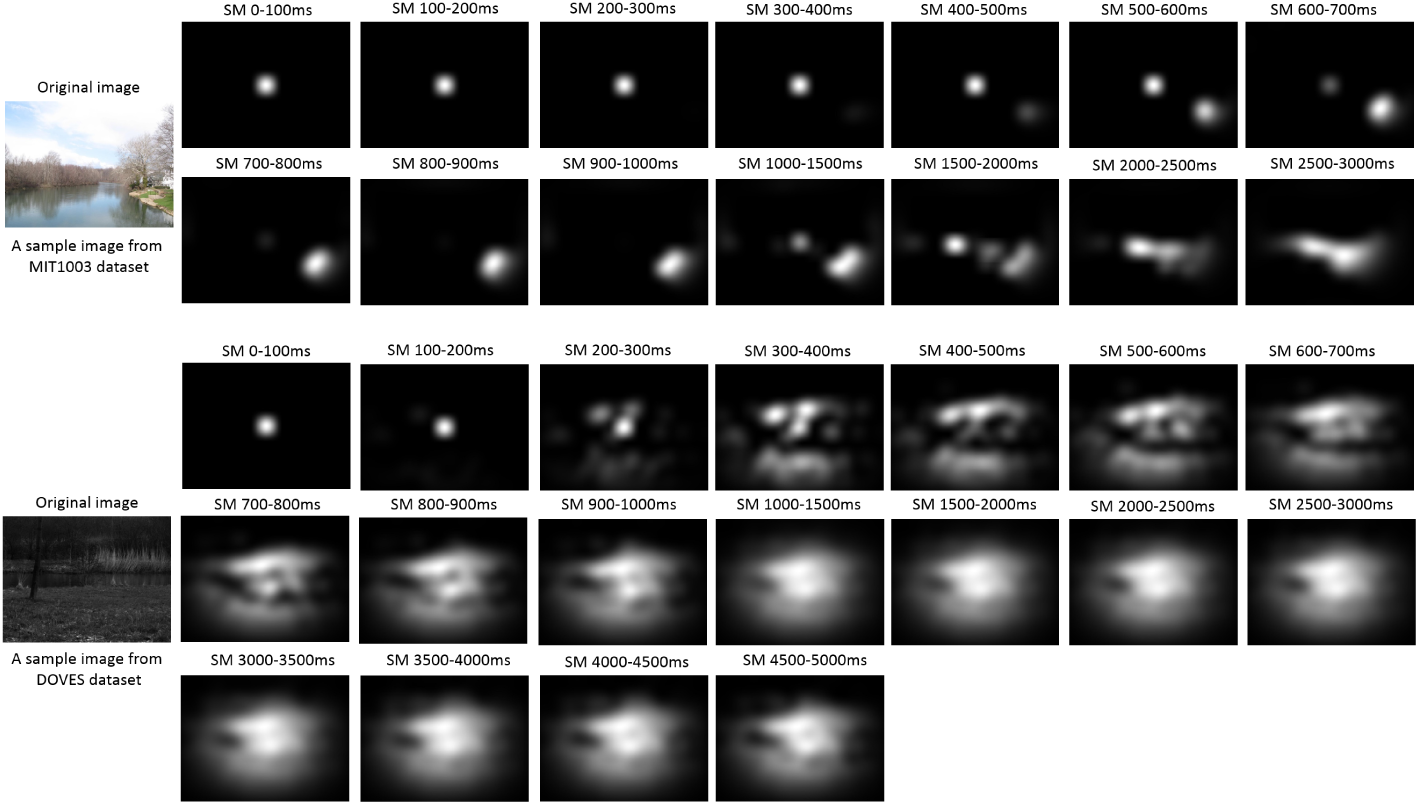
The proposed dynamic saliency map for a sample image in MIT1003 dataset (top) and a sample image in DOVES dataset (bottom).

The NSS, SIM, CC, and KL measures [14] were employed to evaluate the proposed method according to the recommendations given in [67]. SIM, CC, and NSS are similarity measures, and KL is a dissimilarity measure. The results were compared with four bottom-up attention models Itti[9], GBVS[22], Context-Aware[30] and Covsal[29] on MIT1003 dataset and three bottom-up attention models Itti[9], GBVS[22] and Context-Aware[30] on DOVES dataset. Dynamic ground truth for datasets is generated as explained in Section 4. The evaluation results on MIT1003 and DOVES datasets are reported in Figure 4 and Figure 5, respectively. It is worth-noting that the proposed method is a dynamic method and creates a different saliency map for each time interval (SM(t)). Evaluation is done on each time interval separately. We compare the proposed saliency map (SM(t)) for each image to the corresponding ground truth (GT(t)) using the above-mentioned metrics. Finally, the metrics’ average over all images of the dataset is calculated and plotted. In contrast, the competitors are static methods and create only one saliency map (irrespective of time, SM). Therefore, for each image we compare that saliency map to the ground truth extracted for each time interval (GT(t)). Then, the average of evaluation results of all images is calculated and plotted. In the figures, the green line represents the evaluation of ground truth images compared with themselves. As expected, SIM and CC metrics report 1 and KL reports 0. However, NSS metric report different values for each interval.

**Figure 4:**
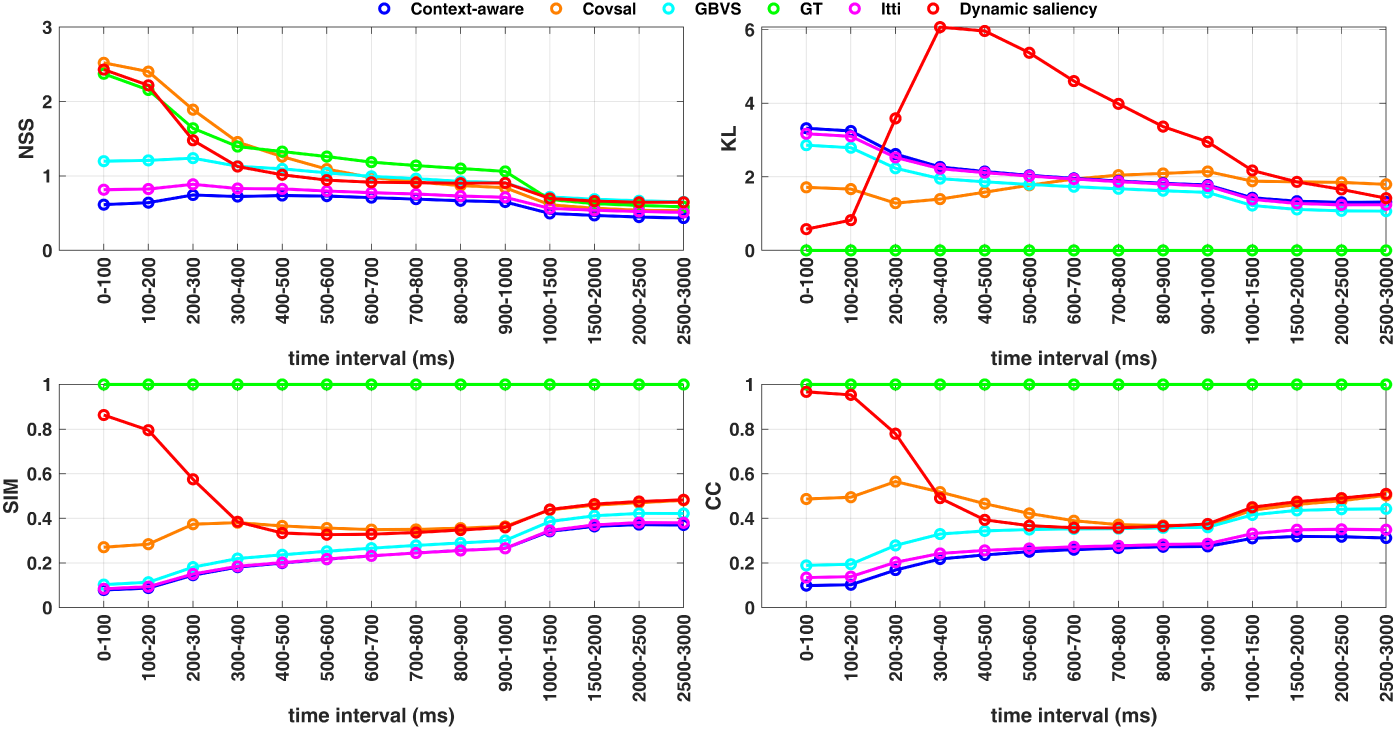
Evaluation results using NSS, CC, SIM, and KL metrics on MIT1003 dataset. Green and red lines indicate the ground-truth and the proposed dynamic saliency(DS) method, respectively. Blue, orange, cyan and magenta indicate our competitors Context-Aware[30], Covsal[29], GBVS[22] and Itti[9], respectively.

**Figure 5:**
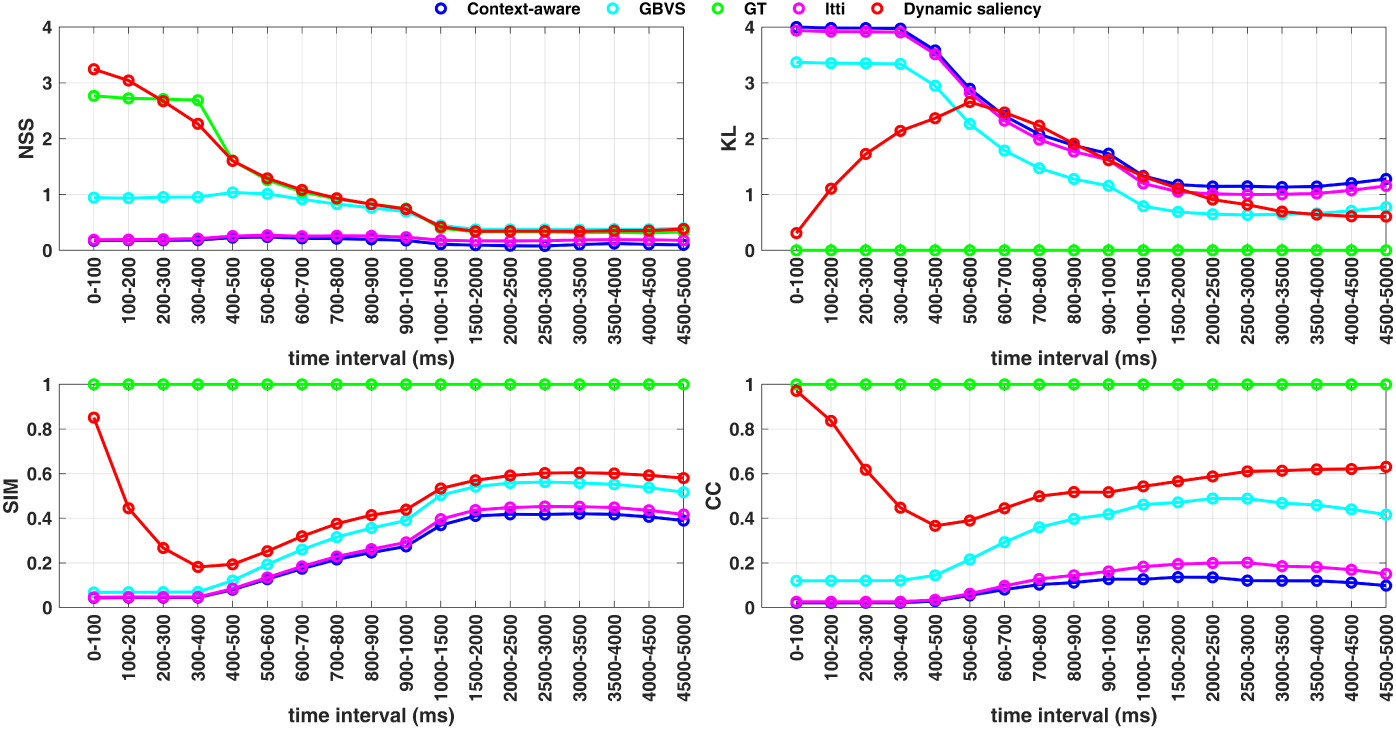
Evaluation results using NSS, CC, SIM, and KL metrics on DOVES dataset. Green and red lines indicate the ground-truth and the proposed dynamic saliency(DS) method, respectively. Blue, cyan and magenta indicate our competitors Context-Aware[30], GBVS[22] and Itti[9], respectively.

The proposed bottom-up saliency detection algorithm is plausible with recent findings about creation of saliency map in V1 [1] or superior colliculus [68]. In both cases, low-level features including color and orientation which are extracted in the retina and V1 are considered vital. Since realizing the bottom-up saliency is fast [69, 70, 6], the first eye movements reflect this type of attention. According to Figure 4, the proposed method has an acceptable performance within 0-400ms after stimulus onset. As the time passes by, top-down attention overrules and starts to compete. Our method is based on low-level features and is developed to predict bottom-up saliency. To generalize the algorithm to be able to anticipate top-down saliency more accurately, a top-down module should be added to the current algorithm. In addition, a proper method for modeling the competition between bottom-up and top-down attention should be included to cover variation in saliency map through time. Figure 5 shows that the proposed method works well on DOVES dataset almost in the entire 0-5 seconds. This dataset is considered a bottom-up dataset compared to MIT1003 as it focuses on low-level features. It explains the better performance of our purely bottom-up saliency detection method on DOVES compared to MIT1003.

It is worth-mentioning that an ideal dynamic saliency prediction algorithm should suggest a saliency map at each moment. However, existing datasets impose the limitation of considering time intervals. We were obliged to define at least 100ms time intervals to make sure that there are enough reliable eye data to generate the ground truth. Increasing the number of subjects in the free-viewing task can help to decrease the length of each time interval to 50ms or less and have a near real-time ground truth. This simplifies evaluation of dynamic saliency detection algorithms.

Finally, we suggest using different strategies for bottom-up and top-down saliency prediction. The former is an unsupervised and built-in mechanism. However, the latter is learning-based. Benefiting from both strategies helps to develop a unified saliency detection algorithm which is able to predict saliency through time. Such algorithm is more biologically-plausible and demanding in computer vision era and applications such as human-computer interaction.

## 6. Conclusion

In this paper, a brain-inspired dynamic bottom-up saliency detection algorithm is proposed inspired by the theory of “A Saliency map in primary visual cortex.” In this method, variations in the low-level features such as color and orientation are extracted and combined within different windows similar to how V1 neurons encode visual information within their RFs. The center of mass of the salient parts of an image is also included in the saliency map. The predicted saliency map is generated by taking pixelwise maximum of weighted versions of the low-level saliency and the center of mass. A thresholding method with a time-dependent threshold is applied to the map beforehand. The weights are adjusted through time to make the prediction dynamic and model how salient areas varies. We evaluated this dynamic saliency detection algorithm using dynamic ground truth generated from MIT1003 and DOVES datasets. Results have shown that the proposed method has an acceptable performance in predicting bottom-up saliency through time.

## Acknowledgment

This work was partially funded by Iran National Science Foundation (grant number 97024773).

## Compliance with Ethical Standards

### Funding

This work was partially funded by Iran National Science Foundation (grant number 97024773).

### Ethical approval

This article does not contain any studies with human participants performed by any of the authors.

## Footnotes

1 http://people.csail.mit.edu/tjudd/WherePeopleLook/index.html

## Notes

### Competing Interest Statement

The authors have declared no competing interest.

